# Comprehensive transcriptomic analysis shows disturbed calcium homeostasis and deregulation of T lymphocyte apoptosis in inclusion body myositis

**DOI:** 10.1101/2021.06.30.450477

**Authors:** Mridul Johari, Anna Vihola, Johanna Palmio, Manu Jokela, Per Harald Jonson, Jaakko Sarparanta, Sanna Huovinen, Marco Savarese, Peter Hackman, Bjarne Udd

## Abstract

**Objective:** Inclusion body myositis (IBM) has an unclear molecular etiology due to the co-existence of characteristic cytotoxic T-cell activity and degeneration of muscle fibers. Using in-depth gene expression and splicing studies, we aimed at understanding the different components of the molecular pathomechanisms in IBM.

**Methods:** We performed RNA-seq on RNA extracted from skeletal muscle biopsies of clinically and histopathologically defined IBM (n=24), tibial muscular dystrophy (n=6), and histopathologically normal group (n=9). In a comprehensive transcriptomics analysis, we analyzed the differential gene expression, differential splicing and exon usage, downstream pathway analysis, and the interplay between coding and non-coding RNAs (micro RNAs and long non-coding RNAs).

**Results:** We observe dysregulation of genes involved in calcium homeostasis, particularly affecting the T-cell activity and regulation, causing disturbed Ca^2+^ induced apoptotic pathway of T cells in IBM muscles. Additionally, LCK/p56, which is an essential gene in regulating the fate of T-cell apoptosis, shows altered expression and splicing usage in IBM muscles

**Interpretation:** Our analysis provides a novel understanding of the molecular mechanisms in IBM by showing a detailed dysregulation of genes involved in calcium homeostasis and its effect on T-cell functioning in IBM muscles. Loss of T-cell regulation is hypothesized to be involved in the consistent observation of no response to immune therapies in IBM patients. Our results show that loss of apoptotic control of cytotoxic T cells could indeed be one component of their abnormal cytolytic activity in IBM muscles.

## Introduction

Inclusion body myositis (IBM) is a late-onset, acquired muscle disease with unclear etiology, and the poorly understood molecular pathogenesis is under debate due to several factors. The CD8+ T-cell infiltration and overexpression of class I MHC antigens in all muscle fibers indicate an autoimmune cascade and are, in fact, the most consistent finding together with the degeneration of myofibers. However, IBM largely remains refractory to immunosuppressive drugs [1], and comprehensive clinical trials have generally been ineffective [2]. A partial clinical and histopathological overlap with other rimmed-vacuolar (RV) myopathies [3] including accumulations of similar proteins in the RVs [4] support a degenerative pathophysiology. Accumulation/aggregation of these misfolded proteins suggests that IBM could be a protein aggregate disease with immune-mediated cytotoxic inflammation as a resulting secondary feature [5]. However, there is a significant variance in nature and the number of accumulated proteins observed in the IBM muscle biopsies [6]. Similar aggregates observed in HIV-associated IBM [7] suggest that protein aggregation can still be a downstream effect of immune dysfunction. Additionally, the occurrence of rare familial cases [8] and a strong association with immune MHC locus 8.1 ancestral haplotype [9, 10] support a possible genetic predisposition for IBM.

Analysis of tissue-specific mRNAs and subsequent RNA-seq based transcriptomics studies focused on understanding the expression of genes, participating pathways, and networks can increase our understanding of underlying pathomechanisms. Prior studies have investigated the differential gene expression in IBM muscles for both the inflammatory and the degenerative pathology [11-17]. However, no study has attempted a comprehensive analysis of RNA-seq data combining differential gene expression, differential exon, and splicing usage along with an in-depth analysis of the relation between dysregulation of coding and regulatory RNAs in IBM muscles.

Our study used RNA extracted from muscle biopsies of IBM patients, of non-myositis RV-myopathy disease group, and a histopathologically group. We first studied the differential expression of coding, long non-coding RNAs (lncRNAs), and micro RNAs (miRNAs) and then evaluated their possible interplay. Additionally, we studied the transcriptome-wide differential exon and splicing usage. We observed a significant association with genes involved in various calcium-related pathways and identified disturbed calcium regulation specific to T cells in IBM muscles, highlighting the relevance of calcium homeostasis for T-cell activity in IBM muscles. In particular, we identified calcium-induced T lymphocyte apoptosis to be disturbed in IBM muscles.

## Materials and methods

### Patients and skeletal muscle biopsies

Muscle biopsies (predominantly Tibialis anterior or Vastus lateralis) from 24 Finnish patients diagnosed with clinically and pathologically defined IBM according to the ENMC criteria [18] were included. The age of onset was 60 ± 11 years (median ± SD), and the age at muscle biopsy was 70 ± 9 years. Additionally, muscle biopsies from six patients with genetically diagnosed Tibial muscular dystrophy (TMD, caused by heterozygous FINmaj mutation the titin gene) [19] were included. In the TMD cohort, the age of onset was 49 ± 11 years, and age at biopsy 54 ± 14 years. Nine muscle biopsies from individuals that underwent leg amputation for reasons other than a muscle disease [20] were also included. These nine biopsies did not show pathologically defined muscle degeneration or inflammation. Age at sampling for amputees was 70 ± 11 years. All muscle biopsies were snap-frozen and stored at −80 **°**C. Muscle biopsies were collected at the Tampere Neuromuscular Research Center, Tampere University Hospital, Finland.

### RNA extraction, selection, and library preparation

Muscle tissue homogenization steps were performed using SpeedMill PLUS (Analytik Jena AG, Germany). RNA was extracted with Qiagen RNeasy Plus Universal Mini Kit (Qiagen, Hilden, Germany) and treated with Invitrogen TURBO DNAse buffer (ThermoFisher Scientific, MA, USA) according to the manufacturers’ instructions. RNA was quantified and qualitatively assessed using High Sensitivity RNA ScreenTape (Agilent Technologies, CA, USA) on Agilent 4200 TapeStation system (Agilent Technologies).

Library preparations and sequencing were performed at Oxford Genomics Center, University of Oxford. For PolyA+ RNA selection, the NEBNext Ultra II Directional RNA Library Prep kit (E7760) for Illumina (NEB, Beverly, MA, USA) was used to prepare strand-specific RNA-seq libraries. Libraries were multiplexed and sequenced on HiSeq4000: 75bp paired-end sequencing (Illumina, CA, USA), and an average of ∼47 million reads per sample were produced. Samples with enough RNA were used for library preparation for small RNA (< 200 nt) selection (18 IBM, nine amputees, and four TMD). NEBNext Small RNA Library Prep Set (E7330) for Illumina was used per the manufacturer’s instructions (NEB). Libraries were multiplexed and sequenced on HiSeq2500: 50bp single-end sequencing (Illumina), and an average of ∼10 million reads per sample were produced.

### RNA-seq data pre-processing, QC, and alignment

Adapter sequences and low-quality bases were removed with fastp [21]. Trimmed sequences were then mapped with STAR 2.7.0d [22] (STAR, RRID: SCR_004463) with index generated from Gencode.v29 human reference (release date 05.2018, based on ENSEMBL GRCh38.p12) and comprehensive gene annotation (primary assembly) using the STAR two-pass method according to the guidelines from the ENCODE project for alignment of long RNA (>200 nt) and small RNA (<200 nt) data.

### RNA-seq quantification and differential gene expression analysis

Uniquely mapped fragments were summarized and quantified (referred to as counts) by featureCounts [23] (featureCounts, RRID: SCR_012919) using Gencode.v29 primary comprehensive gene annotation, which lists 58,780 RNAs including 19,969 protein-coding, 16,066 non-coding, and 22,745 other types of RNAs (primary gene expression analysis). Separate quantification of counts for lncRNA (lncRNA analysis) was done using long non-coding RNA gene annotation from Gencode.v29 (a subset of the primary annotation). Quantification of counts for miRNAs (miRNA analysis) in 31 samples was done using miRBase human miRNA annotation (Release 22.1 October 2018) [24]. Differential gene expression analysis was performed with DESeq2 [25] (v1.26.0) (DESeq2, RRID: SCR_015687) in Rstudio (v1.2.5019) (RStudio, RRID: SCR_000432) based on R (v3.6.3) (R Project for Statistical Computing, RRID: SCR_001905). Counts were normalized with variance stabilizing transformation function within DESeq2. A principal component analysis (PCA) was performed on the gene expression data of the IBM samples compared to amputee and TMD groups. Further, pairwise comparisons between cohorts were performed using the Wald test. Log_2_ fold changes (LFC) were shrunk using ‘ashr’ adaptive shrinkage estimation [26], and results were generated with default independent filtering for increasing power. Only genes with LFC values larger than ±1.5 and a Benjamini-Hochberg adjusted p-value of ≤0.01 were considered further. Genes specifically dysregulated in IBM muscles were considered for downstream analysis.

### Pathway analysis

Ingenuity Pathway Analysis (IPA, QIAGEN Inc.) (Ingenuity Pathway Analysis, RRID: SCR_008653) was used for pathway analysis and enrichment analysis of the obtained differential gene expression data. Using Ingenuity Pathways Knowledge Base (Ingenuity Pathways Knowledge Base, RRID: SCR_008117), IPA mapped and annotated genes to the pathways and predicted activation state based on the direction of changes comparing it with the change in the database.

### Differential splicing analysis

To investigate differential usage of exons and splicing, independent of the differential gene expression analysis, we used QoRTS [27] java-based application (v1.3.6) (QoRTs, RRID: SCR_018665) to prepare counts from exons and splice junctions (known and novel) from the aligned data. Downstream analysis of this data was performed using JunctionSeq [28] (v1.16.0) in R. JunctionSeq results produce a q-value (based on FDR) on gene-level analysis, which considers that one or more exon/junction in this gene is differentially used. A conservative q-value threshold of 0.01 was used to select significant observations. IBM-specific differentially expressed genes and differentially spliced genes were compared (Fig. 1). Statistical over-enrichment analysis for Gene ontology terms in categories: Molecular function, biological process, and cellular component, was performed on results obtained from QoRTs/JunctionSeq using clusterProfiler [29] (clusterProfiler, RRID: SCR_016884). Gene sets were compared using UpSet plot [30].

**Fig. 1.**
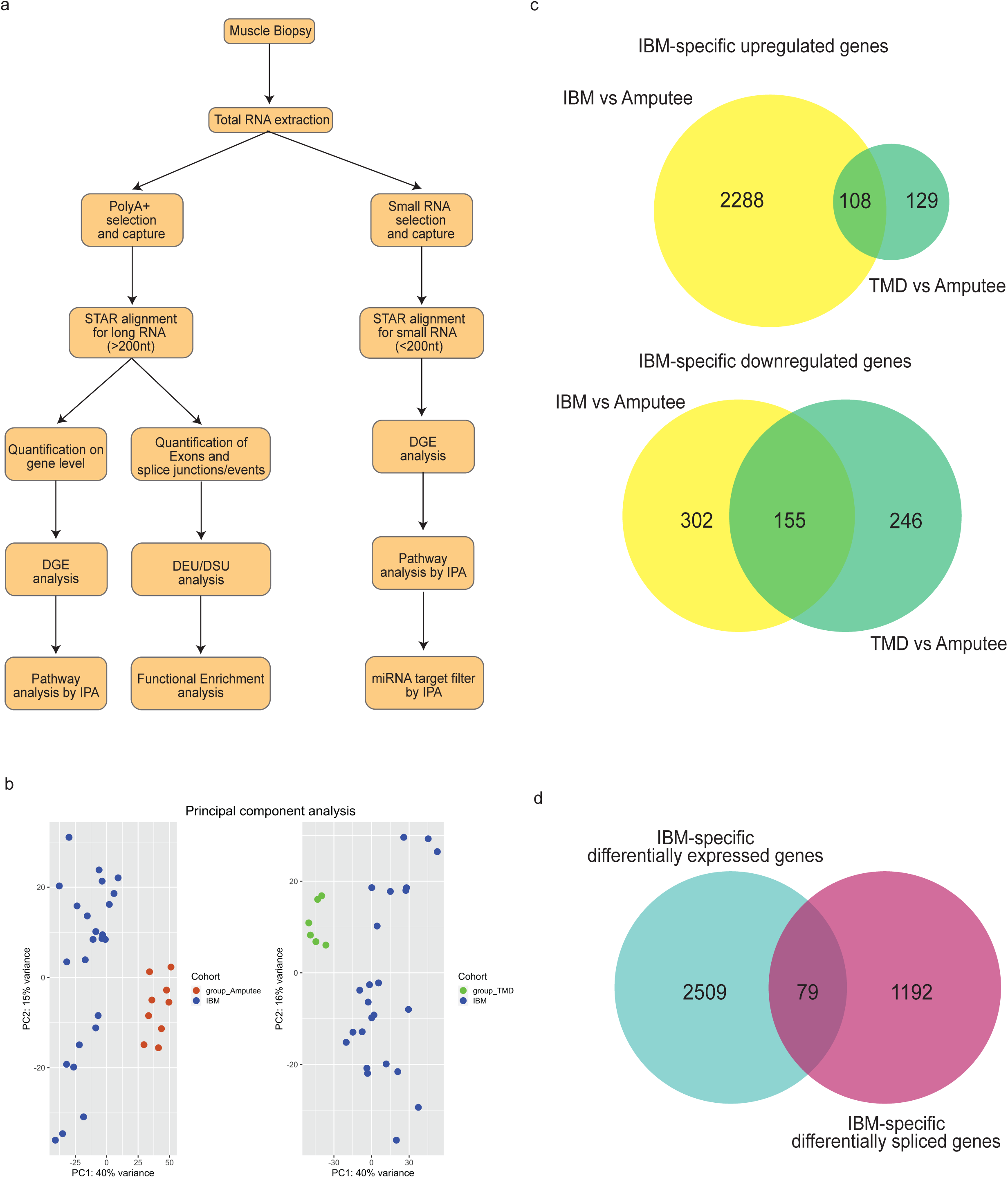
a) Workflow and methodology used in this study. b) Principal component analysis of gene expression results showing the pairwise comparison between different groups: IBM, TMD and Amputees. c) IBM-specific differentially expressed genes were determined by comparing IBM cases with amputee and TMD groups. d) Comparison between IBM specific differentially expressed genes (cyan) and IBM specific differentially spliced genes (magenta).

## Results

### Expression signature in IBM muscles

Fig. 1a shows the summarized workflow of the methodology. The PCA shown in Fig. 1b explains the differences between the three cohorts. Pairwise comparisons were performed to reduce the potential confounding effects of groups, which identified 2,288 and 302 genes specifically up- or down-regulated in the IBM cohort, respectively (Fig. 1c). Non-coding RNA analyses resulted in 497 lncRNAs upregulated, 106 lncRNAs downregulated, 140 miRNAs upregulated, and 126 miRNAs explicitly downregulated in the IBM cohort compared to other groups. These IBM-specific dysregulated RNAs were used for downstream pathway analysis using IPA workflow. The top 15 genes dysregulated specifically in IBM muscles, with their functional annotations and normalized expression in the different cohorts, are shown in Fig. 2.

**Fig. 2.**
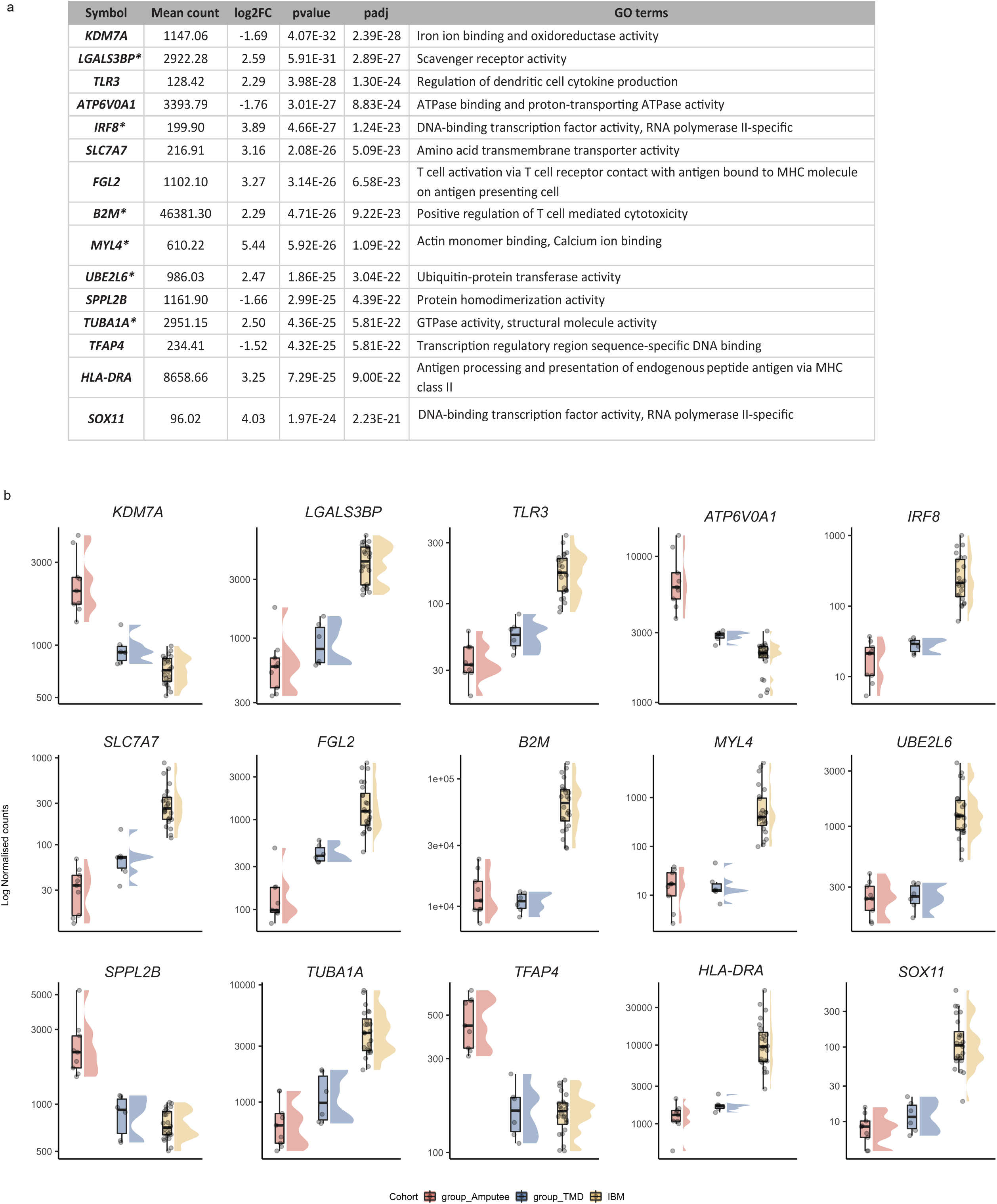
a) Top 15 differentially expressed genes specific to IBM muscles. Log_2_ fold change (log2FC) of IBM versus amputees calculated by DEseq2 after shrinkage estimations. ‘+’/’-’ sign denotes the direction of change, i.e., positive log2FC values indicate overexpressed genes in IBM muscles, and negative log2FC values indicate underexpressed genes in IBM muscles. The p-value of significance and adjusted p-value using the Benjamini-Hochberg corrections and associated GO terms are shown for each gene. Genes marked with * are also observed as significantly dysregulated in Hamann et al. [13] b) Normalized gene expression in the different cohorts is presented as boxplots. Median and quartile values are shown, with whiskers reaching up to 1.5 times the interquartile range. Individual expression levels are shown as jitter points. The raincloud plots illustrate the distribution of data in each cohort. The scaled Y-axis shows log normalized counts.

### Pathway analysis

We performed IPA workflow analysis on IBM-specific dysregulated genes to better understand the pathways and the upstream regulators associated with the observed expression dysregulation. Out of these, 2,588 genes, 596 lncRNAs, and 257 miRNAs mapped to the Ingenuity database. From the primary gene expression analysis, IPA identified 91 pathways as significantly altered. Table 1 shows a summary of the IPA results with the top identified pathways.

**Table 1:** Top 10 dysregulated canonical pathways identified by IPA. The significance of the identified pathway is shown with a p-value and the number of differentially expressed genes observed in the IBM-specific dataset compared to the number of genes present in the database for each pathway.

| Ingenuity Canonical Pathways | p-value | Number of genes in dataset / Number of genes in database |
| --- | --- | --- |
| **Dendritic Cell Maturation** | 5.72E-31 | 106/357 |
| **T Cell Receptor Signaling** | 2.30E-25 | 97/355 |
| **T Cell Exhaustion Signaling Pathway** | 3.12E-25 | 94/338 |
| **Cdc42 Signaling** | 7.63E-24 | 88/315 |
| **iCOS-iCOSL Signaling in T Helper Cells** | 4.10E-23 | 81/280 |
| **CD28 Signaling in T Helper Cells** | 1.13E-22 | 82/290 |
| **OX40 Signaling Pathway** | 4.03E-21 | 68/222 |
| **Calcium-induced T Lymphocyte Apoptosis** | 1.29E-20 | 69/232 |
| **Nur77 Signaling in T Lymphocytes** | 3.07E-20 | 72/253 |
| **Role of NFAT in Regulation of the Immune Response** | 4.57E-19 | 87/360 |

The top upstream regulators in both miRNA and lncRNA analysis are shown in table 2 and table 3, respectively. We identified an increased expression of the lncRNA *DNM3OS* (DNM3 antisense RNA) and *MIAT* (Myocardial infarction associated transcript) from these analyses. IPA suggested this dysregulation may be due to *JDP2* (Jun Dimerization Protein 2) and *TARDBP* (TAR DNA Binding Protein), acting as an upstream regulator of *DNM3OS* and *MIAT* respectively (table 4).

**Table 2:** From the miRNA analysis, upstream binding partners are shown along with their target miRNA. A p-value and associated GO terms are shown.

| Upstream regulator | p-value | GO terms and annotations |
| --- | --- | --- |
| ***AGO2*** | 7.89E-23 | RNA polymerase II complex binding |
| ***SSB*** | 4.74E-19 | RNA binding |
| ***TP53*** | 7.59E-09 | Transcription regulatory region sequence-specific DNA binding |
| **RNA polymerase III** | 4.09E-06 | Synthesis of small RNA, RNA polymerase activity |

**Table 3:** From the long non-coding RNA analysis, upstream binding partners are shown along with their target lncRNA. A p-value and associated GO terms are shown

| Upstream regulator | Target molecule in dataset | p-value | GO terms and annotations |
| --- | --- | --- | --- |
| *JDP2* | *DNM3OS* | 4.10E-03 | DNA-binding transcription factor activity, RNA polymerase II-specific |
| **miR-338-3p** | *NR2F1-AS1* | 4.10E-03 | Negative regulation of gene expression; negative regulation of IL-6 production; negative regulation of cytokine production involved in inflammatory response |
| **miR-150-5p** | *MIAT* | 4.10E-03 | mRNA binding involved in posttranscriptional gene silencing |
| *TARDBP* | *MIAT* | 2.03E-02 | RNA polymerase II cis-regulatory region sequence-specific DNA binding |
| **mir-150** | *MIAT* | 2.63E-02 | mRNA binding involved in posttranscriptional gene silencing |
| *PGF* | *DNM3OS* | 3.43E-02 | Protein binding, signal transduction |
| *FUS* | *RMRP* | 3.43E-02 | mRNA binding, mRNA stabilization |
| *DDX58* | *EGOT* | 4.61E-02 | double-stranded RNA binding |

**Table 4:**
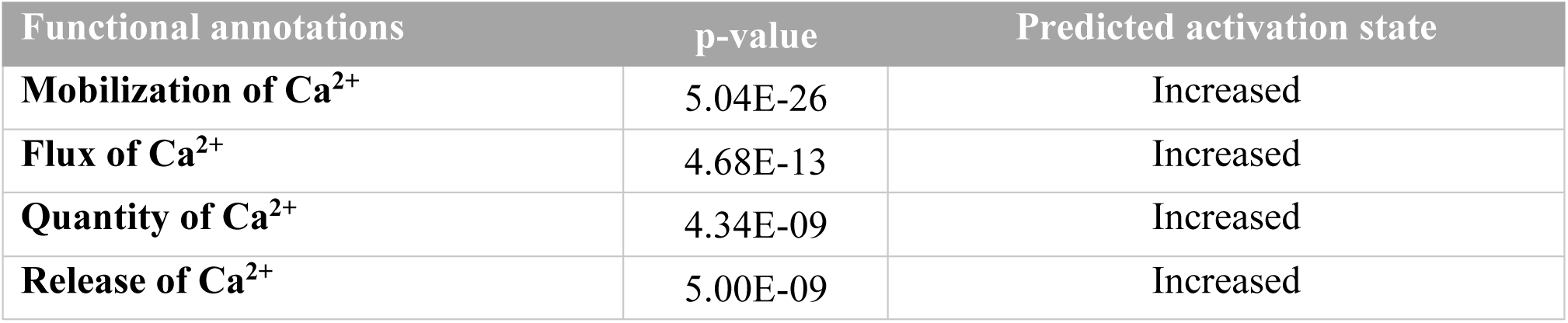
In cell signaling processes, different pathways associated with calcium homeostasis are shown along with their p-value and a prediction state.

| Functional annotations | p-value | Predicted activation state |
| --- | --- | --- |
| **Mobilization of Ca<sup>2+</sup>** | 5.04E-26 | Increased |
| **Flux of Ca<sup>2+</sup>** | 4.68E-13 | Increased |
| **Quantity of Ca<sup>2+</sup>** | 4.34E-09 | Increased |
| **Release of Ca<sup>2+</sup>** | 5.00E-09 | Increased |

### Dysregulation of calcium-related pathways in IBM muscles

IPA identified calcium-induced T lymphocyte apoptosis as one of the most significant pathways dysregulated in IBM muscles (table 1). Our IBM-specific dataset contained 69 genes with significant dysregulation out of the 232 genes annotated in this pathway. A part of this pathway, including the major players, is shown in Fig. 3. Another pathway outside the top results identified that 29 genes (29/208, p = 7.05E-03) significantly dysregulated in our dataset are also involved in calcium signaling. These results prompted us to investigate further for calcium related issues in cellular signaling, and we found that IPA also detects dysregulation of the following processes, mobilization of Ca^2+^ (80 genes), the release of Ca^2+^ (33 genes), quantity of Ca^2+^ (51 genes) and flux of Ca^2+^ (51 genes), as significantly disturbed in IBM muscles (table 4).

**Fig. 3.**
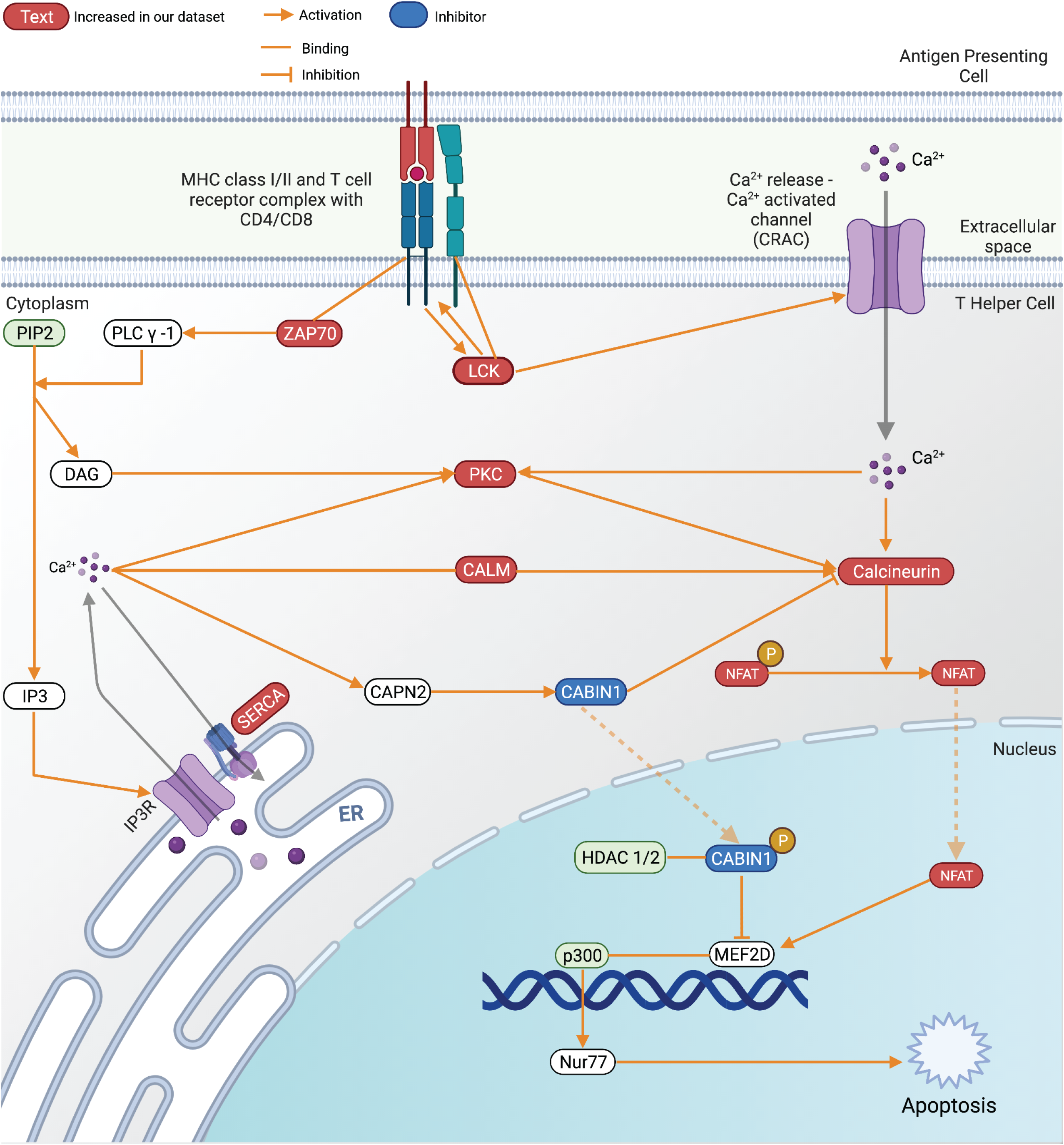
The calcium-induced T lymphocyte apoptosis pathway with gene expression changes observed in IBM compared to groups. Created with BioRender.com

### Altered exon usage and splicing pattern in IBM muscles

To explore IBM-specific exon usage, we performed an independent transcriptome-wide differential splicing analysis in our three cohorts. We obtained a list of 1,271 differentially spliced genes in IBM from our differential splicing analysis. These transcripts either showed IBM-specific increased usage of a known junction or a known exon or contained a novel exon-exon junction resulting in an alternative isoform. To understand the diverse portfolio of mature mRNAs created from pre-mRNAs, we used gene ontology over-enrichment analysis on these 1,271 differentially spliced genes and identified the first splicing signature specific to IBM muscles. To understand the different classes over-represented in these genes, we performed statistical over-enrichment analysis using clusterProfiler for all three GO categories as seen in Fig. 4 a,b,c. Our analysis showed an enrichment of genes involved in the structure and organization of actin filaments assembly in IBM muscles and, interestingly, proteins involved in mRNA processing and metabolism.

**Fig. 4.**
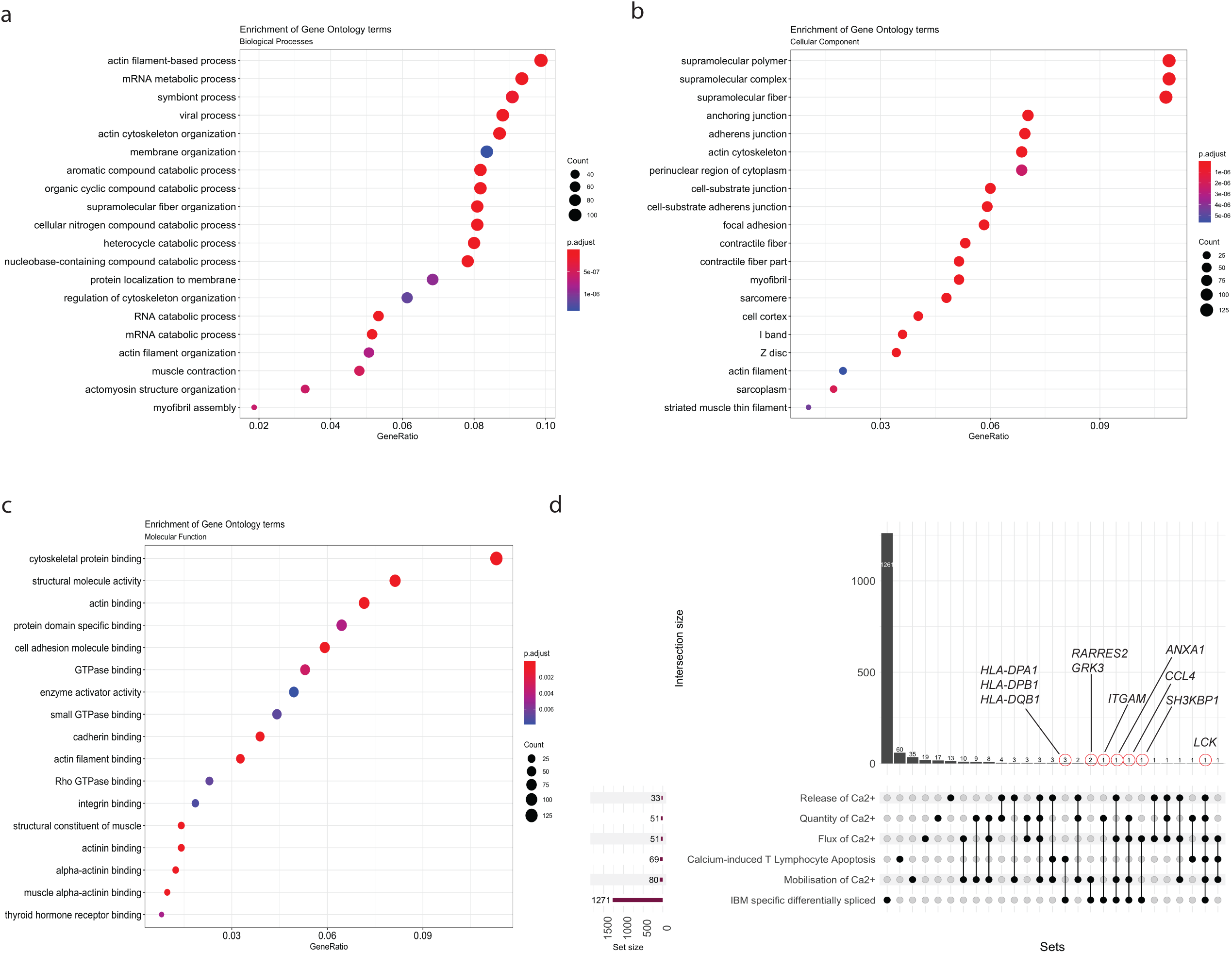
Statistical over-representation tests were performed on a list of differentially spliced RNAs, using clusterProfiler for a) Biological Processes, b) Cellular component, and c) Molecular function. d) An UpSet plot is shown comparing six different sets, namely, IBM specific differentially spliced (1,271 genes), mobilization of Ca^2+^ (80 genes), calcium-induced T lymphocyte apoptosis (69 genes), the flux of Ca^2+^ (51 genes), quantity of Ca^2+^ (51 genes), and release of Ca^2+^ (33 genes). Dots and lines represent subsets of different lists. The horizontal bar graph (wine color) represents the size of each set, while the vertical histogram (black) represents the number of RNAs in each subset. The 10 RNAs that are both differentially expressed and differentially spliced are shown with a red circle with their gene names (black).

We then compared the list of differentially spliced genes with differentially expressed genes in our analysis and found an overlap of 79 genes (Fig. 1d). Next, we wanted to observe the overlap between six different sets of genes, namely IBM specific differentially spliced genes, calcium-induced T Lymphocyte apoptosis, Mobilization of Ca2+, Flux of Ca2+, Quantity of Ca2+, and Release of Ca2+ (Fig. 4d). We observed 10 genes to be associated with calcium-related processes; *HLA-DPA1, HLA-DPB1*, and *HLA-DQB1* are associated with calcium-induced T Lymphocyte apoptosis, *ANXA1* is associated with mobilization, flux, and release of Ca^2+^, *CCL4* is associated with mobilization, flux, and quantity of Ca^2+^, *GRK3* and *RARRES2* are associated with mobilization, *SH3KBP1* with flux, and *ITGAM* with the quantity of Ca^2+^. In particular, one specific differentially spliced gene, *LCK*, is part of all six sets.

Fig. 5a shows the gene expression of *LCK* in three cohorts, with expression in IBM muscles being significantly higher than the others (log_2_FC = +2.86, padj=3.50E-11, ranking = 355/2590). Additionally, Fig. 5b shows the differential splicing pattern observed in *LCK* in all three groups. The highlighted E016 corresponds to an alternative exon (chr1:32274818-32274992, GRCh38).

**Fig. 5.**
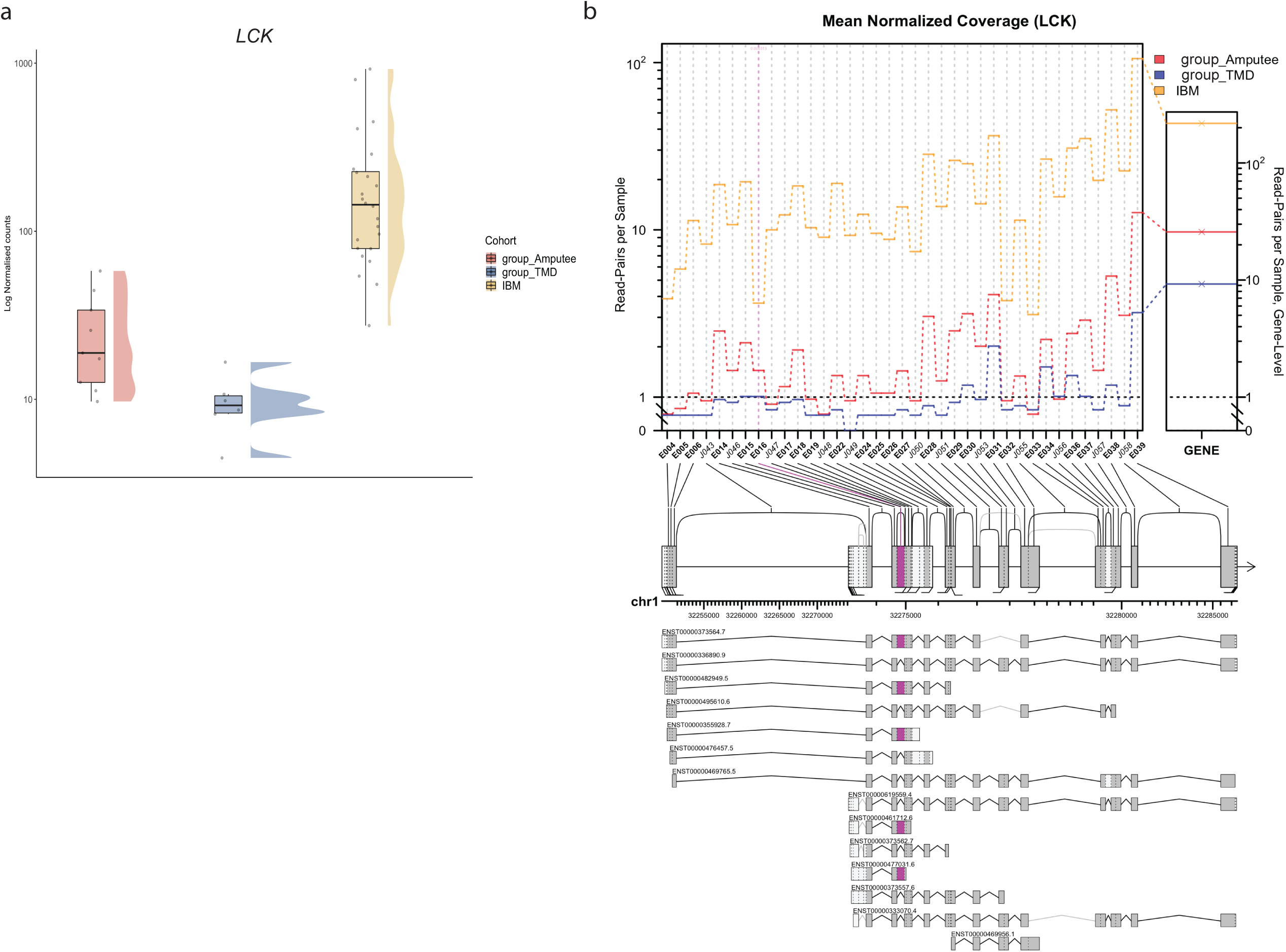
a) Normalized LCK expression in the different cohorts (as explained in Fig. 2b). b) Altered isoform expression of *LCK* using JunctionSeq showing estimated normalized mean read-pair count for each exon and splice junctions in the different cohorts (left) as well as for the whole *LCK* gene (right). The significantly alternatively spliced feature, E016 (pink), corresponds to chr1:32274818-32274992 (GRCh38). The alternative *LCK* transcripts used in the JunctionSeq analysis are shown below with their corresponding ENSEMBL identifiers.

## Discussion

In this study, we aimed to identify a more detailed IBM-specific molecular signature, using different RNA-seq based methods that can help us explore the inflammatory and degenerative parts in depth. Antigen-driven T-cell cytotoxicity is the most reproducible and plausible part of the complex molecular pathomechanism in IBM. However, it remains unknown what antigen drives this IBM-specific immune cascade.

As part of the RV pathology, accumulated proteins or the unfolded protein response have been hypothesized to prompt an immune reaction [5]. A recent unbiased proteomics study dissected these RVs in IBM [31]. Interestingly, the protein encoded by one of our top differentially expressed genes, *MYL4*, is also detected in the RVs in IBM along with *ANXA1*, which is both differentially expressed and differentially spliced in IBM muscles. In our study design, we considered TMD, another RV muscle disease but without immune involvement, to understand if there are any RV-specific antigens in IBM muscles. Additionally, using age-matched histopathologically normal muscles from amputees, we aimed to understand if general inflammatory signatures can be replicated and studied in more detail using additional methods such as non-coding RNAs and differential splicing studies. Consequently, our strong study design and robust methodology helped us replicate findings from previous studies [11-17] and identify essentially new calcium-related issues in IBM muscles and their link with the altered T-cell cytotoxicity in IBM muscle fibers.

We found that several genes contributing to calcium homeostasis are differentially expressed in IBM muscles resulting in dysregulation of several critical pathways, specifically, calcium-induced T lymphocyte apoptosis and related Nur77 signaling. Ca^2+^ is a universal second messenger in T cells, and it is known to regulate proliferation and differentiation of T cells and T-cell effector functions [32]. The complexity and duration of Ca^2+^ signals and resultant cytoskeletal rearrangements determine the fate of T cells in response to an antigen [33]. On one hand, a short-term increase in intracellular Ca^2+^ concentration results in the cytolytic activity of T cells; on the other hand, prolonged elevation results in proliferation, differentiation, and maturation of näive T cells into Th1, Th2, and Th17 subtypes and the production of cytokines[32].

Ca^2+^ signaling is known to optimize the interaction between T cells and antigen-presenting cells [33]. The binding of antigen/MHC complexes (CD8^+^-MHC class I/CD4^+^-MHC class II) to T-cell receptors (TCR) activates Src-family protein tyrosine kinases, e.g., LCK and FYN at the cytoplasmic side of the TCR/CD3 complex. Additionally, activation of ZAP-70, a tyrosine kinase associate protein, results in the phosphorylation and activation of the intracellular enzyme phospholipase C-γ1 (PLC-γ1) [32, 33]. PLC-γ1 hydrolyses phosphatidylinositol 4,5-biphosphate (PIP2) to produce two other second messengers, inositol 1,4,5-triphosphate (IP3) and diacylglycerol (DAG). IP3 binds to its receptor (IP3R) on the endoplasmic reticulum (ER) membrane to promote rapid release of Ca^2+^ from ER to the cytosol [32]. However, this release of Ca^2+^ is insufficient for antigen-derived T-cell fate but results in depletion of intracellular Ca^2+^ triggering a rapid influx of Ca^2+^ through activation and opening of Ca^2+^ release-Ca^2+^ activated channels (CRAC) on the plasma membrane formed by different STIM1/ORAI1 combinations [34]. The duration of Ca^2+^ influx is vital for activating the calcineurin-dependent nuclear factor activate T cells (NFAT) transcription pathway [32]. In the cytoplasm, calcineurin removes excess phosphate residues from the N terminus of NFAT, promoting its entry into the nucleus. Disruptions in NFAT signaling can cause several phenotypes, including cardiovascular, musculoskeletal, and immunological diseases [35]. Meanwhile, DAG, another secondary messenger, activates protein kinase C (PKC), which in turn activates the nuclear factor kappa B (NFκB). The duration and complexity of Ca^2+^ signals drive the NFAT/NFκB signaling and determine downstream T-cell activation.

The genes in the NR4A family (NR4A1/Nur77, NR4A2/Nurr1, NR4A3/Nor1) act as critical molecular switches in cell survival and inflammation. Human *NR4A1* encodes for a homolog of a mouse protein called Nur77, a zinc transcription factor expressed as an early gene in T cells upon antigen-TCR interaction. In addition to being a transcriptional activator, Nur77 has an apoptotic role in T regulatory fate [36] and other non-genomic proapoptotic functions via mitochondrial interactions with Bcl-2 [37]. T cells deficient in Nur77 have been shown to have high proliferation, enhanced T-cell activation, and increased susceptibility for T-cell-mediated inflammatory diseases [38]. The expression of Nur77 is Ca^2+^ dependent and is controlled by the myocyte enhancer factor 2 (MEF2) transcription factor [39], whose DNA-binding and transcriptional activity is enhanced by Calcineurin. Another calcium-dependent transcription factor, CABIN1, acts as a transcriptional repressor of MEF2, thus keeping the Nur77 promoter silent in the absence of a TCR signal [40]. The interaction between CABIN1 and calcineurin is influenced by intracellular Ca^2+^ and PKC activation, resulting in hyperphosphorylation of CABIN1 and its subsequent transcription repressing activity. An increase in intracellular Ca^2+^ concentration activates the interaction of the calmodulin family of genes (CALM) with CABIN1, triggering the dissociation of MEF2 from Cabin and MEF2 to become transcriptionally active [41]. In the nucleus, NFAT interacts with MEF2 and enhances its transcriptional activity by recruiting the co-activator p300 for the transcription of Nur77.

In our dataset, 69 genes mapping to the calcium-induced T Lymphocyte apoptosis and 72 genes mapping the Nur77 signaling in T Lymphocytes are differentially expressed in IBM muscles. As seen in Fig. 3, several essential genes like *ZAP70, LCK*, different subunits of Protein Kinase C, and *ATP2A1* which encodes for SERCA, are significantly changed in IBM muscles. Additionally, we also observed genes associated with the mobilization of Ca^2+^, the release of Ca^2+^, the quantity of Ca^2+,^ and the flux of Ca^2+^ as significantly dysregulated in IBM muscles, indicating a possible widespread disturbance with the handling of calcium entry and release in cells. In T cells, especially, this disturbance could dramatically impact their activation, differentiation, and most likely, the regulation of T-cell apoptosis will be disturbed.

Apoptosis in T cells is necessary to resolve their inflammatory activity, and defective or delayed apoptosis may contribute to the pathogenesis of inflammatory diseases [42]. In this scenario, loss of apoptotic control could be one mechanism explaining the lack of immune-suppressive therapeutic effect in IBM [43].

The diversity of the skeletal muscle proteome is, among others, dependent on the diversity of exon usage in pre-mRNAs [44]. From our transcriptome-wide splicing analysis within the differentially expressed genes, we identified 79 genes, out of which ten are associated with different calcium-related functions. Amongst these, LCK is a T lymphocyte-specific protein tyrosine kinase involved in downstream events of antigen–TCR interaction. LCK/p56 is essential in transducing signals leading to apoptotic cell death in mature T cells [45], and its activity is tightly regulated to protect against hyperactivation of T cells and autoimmunity, thus maintaining T-cell homeostasis [46]. Moreover, LCK also selectively influences the flux and release of calcium in cells [47]. In our analysis, LCK is both differentially expressed and differentially spliced in IBM muscles. Disturbed T-cell apoptosis and the dysregulation of LCK in IBM muscles provide novel insights into the molecular mechanisms of IBM. Considering the crucial regulatory activity of LCK, it might be a potential therapeutic target for IBM patients.

We also observe dysregulation of several non-coding RNAs in our study. Previously, Hamann and colleagues have discussed lncRNAs in the context of IBM [13]. The benefits of our study design, especially the homogenous molecular pathology and the larger sample size, let us dig deeper into the dysregulation of lncRNAs specific to IBM muscles. We identified that JDP2 (DNA binding transcription factor) and TARDBP/TDP-43 (DNA and RNA binding protein) might have altered regulator activity since their downstream non-coding partners (*DNM3OS* and *MIAT*, respectively) are significantly overexpressed IBM muscles. Additionally, both these proteins are specific to RNA polymerase II (RNA Pol II), facilitating transcription and pre-mRNA maturation. Alteration in RNA or DNA binding proteins (expression or localization) associated with the activity of the spliceosome machinery can directly affect the downstream events. Since TDP-43 is accumulated in RVs, one possibility is that the unavailability of TDP-43 can affect its transcription and splicing activities. The normal expression of *TARDBP* we observe in IBM patients is expected and is in coherence with the previous reports [48]. In inherited muscle diseases, damaging variants in the disease-associated gene can result in mislocalization and accumulation of mutant protein in the muscle fibers. Previous studies have reported rare exonic variants in genes, including *VCP* and *SQSTM1* in IBM [49, 50]. However, in our cohort of IBM patients, there were no rare exonic *TARDBP, VCP*, or *SQSTM1* variants [9] that could suggest a possible association with abnormal protein turnover and accumulation/aggregation. Therefore, further evidence to suggest the pathogenic role of variants in such genes and their downstream effect on pathological protein accumulation in IBM is still missing. However, the potential downside of TDP-43 not being available for its traditional roles, such as effective splicing, because of the aggregation is noteworthy. Further evidence of possible dysregulation of splicing in IBM muscles comes from our differential splicing results where proteins involved in mRNA processing, transcription, and regulation are enriched, suggesting that additional studies are needed to understand the possible impact of dysregulated mRNA processing in IBM muscles.

Previously, Pinal-Fernandez and colleagues observed that calcium-induced T lymphocyte apoptosis was a significant IBM-specific dysregulated pathway in their extensive analysis of different inflammatory myopathies but did not comment further on the possible importance [17]. Additionally, using a smaller sample size, Amici and colleagues identified calcium signaling as one of the significant disturbed pathways in IBM muscles and hypothesized its potential major role [14]. Furthermore, previous gene expression studies in IBM have analyzed data primarily from microarrays [11, 12, 15]. Only recently paired-end reads RNA-seq have been used in IBM studies [13, 14, 16, 17]. While microarray-based analyses are comparable for differential expression studies, RNA-seq based methodologies are superior for in-depth transcriptome analysis. In our study design, we used matched muscle biopsies and state-of-the-art RNA-seq analysis tools. Our analyses show novel molecular events in IBM muscles which increase our understanding of IBM and provide valuable additions to improve the therapeutic interventions considering the disturbed calcium homeostasis, dysregulation of LCK, and associated deregulation of apoptotic control of T cells in IBM muscles.

## Acknowledgments

The authors would like to thank the patients and their families. We thank the Oxford Genomics Centre at the Wellcome Centre for Human Genetics (funded by Wellcome Trust grant reference 203141/Z/16/Z) for the generation and initial processing of the sequencing data. We would like to thank CSC – IT Center for Science, Finland, for its computational resources. We thank Merja Soininen, Talha Qureshi and Eini Penkkimäki for their technical assistance. We would like to thank Peter-Bram ‘t Hoen for his critical review of this manuscript.

## Declarations

## Author Contributions

MJ: Conceptualization of the study, funding acquisition, data analysis and curation, methodology, project administration, visualization, writing the original draft, review, and editing of the manuscript.

AV: data analysis, methodology, writing the original draft.

JP: Patient samples and data collection, writing the original draft.

MJ: Patient samples and data collection, review, and editing of the manuscript.

PH.J.: Conceptualization of the methodology, review, and editing of the manuscript.

JS: Data analysis, review, and editing of the manuscript.

SH: Patient samples and data collection, review, and editing of the manuscript.

MS: Conceptualization of the study and methodology, review and editing of the manuscript.

PH: Funding acquisition, project administration, and review and editing of the manuscript.

BU.: Conceptualization of the study, funding acquisition, supervision, project administration, review, and editing of the manuscript.

## Ethical approval

The study was performed in line with the principles of the Declaration of Helsinki. Ethical approval for this study falls under HUS:195/13/03/00/11. Informed consent from the patients was obtained at the time of sample collection.

## Data availability

Raw counts and normalized DESeq2 counts from polyA+ RNAs and miRNAs are available in GEO as superseries GSE151758.

